# Identification of parthenogenesis-inducing effector proteins in *Wolbachia*

**DOI:** 10.1101/2023.12.01.569668

**Authors:** Laura C Fricke, Amelia RI Lindsey

## Abstract

Bacteria in the genus *Wolbachia* have evolved numerous strategies to manipulate arthropod sex, including the conversion of would-be male offspring to asexually reproducing females. This so-called “parthenogenesis-induction” phenotype can be found in a number of *Wolbachia* strains that infect arthropods with haplodiploid sex determination systems, including parasitoid wasps. Despite the discovery of microbe-mediated parthenogenesis more than 30 years ago, the underlying genetic mechanisms have remained elusive. We used a suite of genomic, computational, and molecular tools to identify and characterize two proteins that are uniquely found in parthenogenesis-inducing *Wolbachia* and have strong signatures of host-associated bacterial effector proteins. These putative parthenogenesis-inducing proteins have structural homology to eukaryotic protein domains including nucleoporins, the key insect sex-determining factor Transformer, and a eukaryotic-like serine-threonine kinase with leucine rich repeats. Furthermore, these proteins significantly impact eukaryotic cell biology in the model, *Saccharomyces cerevisiae*. We suggest these proteins are parthenogenesis-inducing factors and our results indicate this would be made possible by a novel mechanism of bacterial-host interaction.

## INTRODUCTION

The evolution of eukaryotic organisms has been continuously sculpted by relationships with intracellular microbes. The characteristics of these intracellular organisms have been driven by strong selection pressures to manipulate host physiology in favor of their own transmission and persistence (Moran, et al. 2008). Many endosymbionts can be transmitted maternally but not paternally, an asymmetry that can result in sexual and genetic conflicts (Perlman, et al. 2015). This phenomenon is exemplified by one of the most prevalent endosymbionts on earth, the bacterium *Wolbachia*. *Wolbachia* are maternally transmitted alphaproteobacteria (Rickettsiales) in many filarial worms and arthropods, including around half of all insect species (Kaur, et al. 2021). Most remarkably, *Wolbachia* can drastically alter host reproductive outcomes in favor of their own transmission and spread, including the conversion of would-be male offspring to *Wolbachia*-transmitting genetic females: i.e., the induction of parthenogenesis.

Parthenogenesis induction (PI) is arguably the most advantageous reproductive phenotype for *Wolbachia*, as it is the only known symbiont-mediated reproductive manipulation in which host populations can be sustained with females only, resulting in all hosts being avenues for transmission of *Wolbachia* to the next generation. To date, PI *Wolbachia* strains have been confirmed in a range of haplodiploid arthropods, including parasitoid wasps, thrips, and mites (Ma and Schwander 2017). Haplodiploidy is a sex determination system that relies on ploidy of the embryo: in the simplest form, males are haploid and females are diploid. In many lineages this difference in ploidy derives from whether a mother fertilizes a given egg, thus restoring diploidy (Gietz and Woods 2002) (Figure 1A). However, in the presence of PI-*Wolbachia*, diploid females can be produced from unfertilized eggs through a fertilization-independent: *Wolbachia*-mediated restoration of diploidy (Figure 1B).

**Figure 1.**
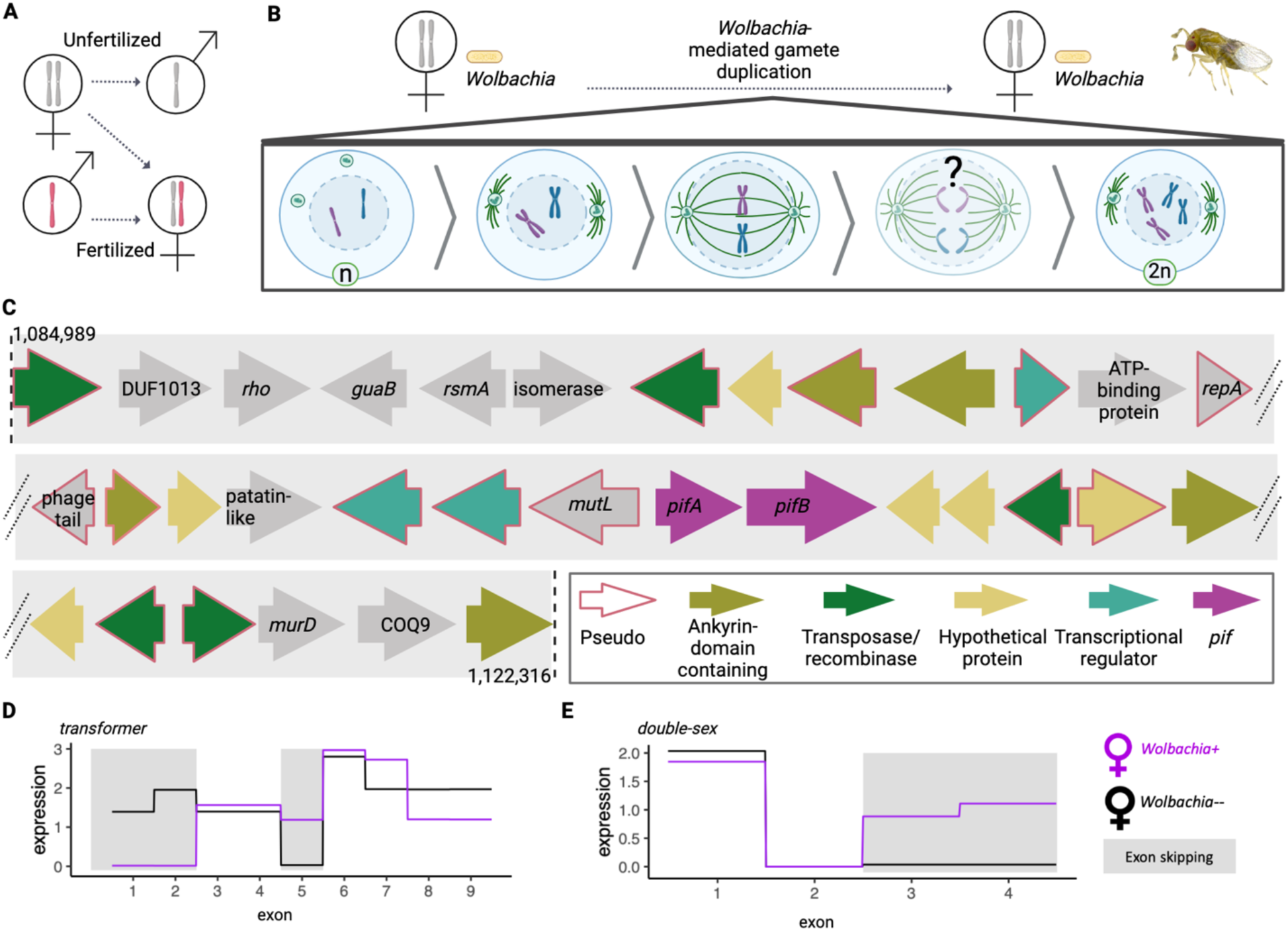
*Wolbachia-mediated parthenogenesis* and putative PI factors. (**A**) Haplodiploid sex determination in the absence of *Wolbachia*. Haploid males develop from unfertilized eggs, and diploid females develop from fertilized eggs. **(B)** Parthenogenesis-inducing *w*Tpre *Wolbachia* causes gamete duplication via a failed anaphase during the first mitotic division in *Trichogramma* embryos. **(C)** Candidate PI factors (*pifs*) are located within a degraded eukaryotic association module. The schematic shows coding regions from base pairs 1,084,989 to 1,122,316 of the *w*Tpre reference (NCBI Reference Sequence: NZ_CM003641.1 (Lindsey *et al*, 2016)). Named loci are in gray, others are colored by annotation according to the legend. Pseudogenized loci (“Pseudo”) are indicated with a pink colored border. **(D)** Exon use of *transformer* and **(E)** *double-sex* in female *Trichogramma pretiosum* with and without *Wolbachia* infections. Exons that are differentially skipped between *Wolbachia* infected and uninfected wasps are indicated with grey boxes.

PI has evolved multiple times across *Wolbachia* as well as in several other genera of arthropod endosymbionts (Ma and Schwander 2017). Different occurrences of microbe-mediated PI are hypothesized to be made possible through different mechanisms, as inferred by cytogenetic approaches combined with estimates of heterozygosity (Stouthamer and Kazmer 1994a; Gottlieb and Zchori-Fein 2001; Weeks and Breeuwer 2001; Pannebakker, et al. 2004; Ma and Schwander 2017). For example, some PI-*Wolbachia* target meiosis to generate diploid gametes, while others target mitosis to convert haploid embryos to diploid. The PI-*Wolbachia* strains *w*Tpre and *w*Lcla cause a failed anaphase in the first mitotic division of the embryo leading to a single diploid nucleus in the parasitoid *Trichogramma* (Figure 1B) (Stouthamer and Kazmer 1994a; Pannebakker, et al. 2004), while the strain “*w*Uni” causes gamete duplication through a tetraploid second mitotic prophase in a different parasitoid genus (Gottlieb and Zchori-Fein 2001), and “*w*Bpra” induces functional apomixis in a species of mite (Weeks and Breeuwer 2001). Thus, PI bacteria have developed a variety of mechanisms for impacting host chromosome segregation.

In addition to differences in the timing and mechanisms for altering ploidy, there appears to be variation in whether a change in ploidy is sufficient to facilitate female development. In several PI systems, diploidization of embryos appears to be necessary but not sufficient, as evidenced by the occasional presence of diploid phenotypic males, and even diploid individuals with intersex characteristics, especially when *Wolbachia* titers have been experimentally reduced (Tulgetske 2010; Lindsey and Stouthamer 2017). Incomplete sexual differentiation to a diploid, fully morphological female implies a two-step mechanism of parthenogenesis induction, wherein both diploidization and impacts on female-specific differentiation are required (Ma, et al. 2015).

Mounting evidence suggests that a number of PI-*Wolbachia* strains manipulate the host’s gene expression cascade that is responsible for female development (Sugimoto and Ishikawa 2012; Geuverink, et al. 2018). Across Hymenoptera, this sexual differentiation pathway centers around the gene *transformer*, the primary signal for female development (Verhulst, et al. 2010). The sex determination cascade involves alternative splicing of *transformer* transcripts to exclude a premature stop codon to produce a functional, female-specific Tra protein, TraF. Male-specific splicing yields a nonfunctional “TraM” protein resulting in male-specific splicing of downstream transcripts, and therefore male development. In the parasitoid wasp *Leptopilina clavipes*, wasps infected with *Wolbachia* strain “*w*Lcla” exhibit differential splicing compared to uninfected lines of the same species, suggesting an interaction with the sex determination system, which could be the key to ensuring female development in such a two-step PI system (Wu, et al. 2020).

Despite the discovery of microbe-mediated transitions to asexuality more than 30 years ago (Stouthamer, et al. 1990), the precise cellular mechanisms and bacterial genes responsible for manipulating host mitosis and reproductive biology have remained elusive. This is largely due to a paucity of genetic tools for *Wolbachia*, and, because PI-*Wolbachia* are not known to occur in genetically tractable models such as *Drosophila*, or even the model parasitoid *Nasonia.* Here, we identify two genes that are unique to “two-step” PI *Wolbachia*: *w*Lcla infecting *Leptopilina clavipes*, and *w*Tpre infecting *Trichogramma pretiosum* (Stouthamer and Kazmer 1994a; Pannebakker, et al. 2004).We show that these two proteins resemble bacterial effector proteins and that they significantly impact eukaryotic biology. Most notably, one of these proteins has strong structural homology to the insect sex determining factor, *transformer.* Together, these proteins are strong candidates for parthenogenesis-inducing effector proteins.

## METHODS

### Genomics, domain prediction, and protein modeling

Thirty-three complete or near-complete *Wolbachia* genome assemblies representing nine Supergroups (A, B, C, D, E, F, J, L, T) were used in comparative genomic analyses to identify putative parthenogenesis-inducing factors (Supplemental Table 1). We specifically included the two PI *Wolbachia* strains with a shared cytological mechanism for PI, *w*Tpre and *w*Lcla that infect the parasitoid wasps *Trichogramma pretiosum* and *Leptopilina clavipes*, respectively (Stouthamer and Kazmer, 1994; (Pannebakker, et al. 2004). Previously published RNA-seq data (Wu, et al. 2020). were queried to visualize exon usage of *transformer* and *double-sex* transcripts in genetically matched female *Trichogramma pretiosum* with and without their *Wolbachia* infections (asexual and sexual, respectively). All genome sequences and PGAP “RefSeq” annotations (Tatusova, et al. 2016) were used to build orthologous groups of *Wolbachia* proteins using ProteinOrtho v5.15 with default parameters (Lechner, et al. 2011). We identified groups of orthologous genes that were unique to the two PI-*Wolbachia*. Protein domain prediction was performed with HHpred (Söding, et al. 2005; Zimmermann, et al. 2018) using SMART_v6.0, PDB_mmCIF70_12_Oct, and Pfam-A_v34 as reference databases. Significant matches to structural domains were determined as those with probabilities greater than 50%, or, greater than 30% and in the top five hits, as per published HHpred best practices (Söding, et al. 2005; Zimmermann, et al. 2018). 3D protein modeling was performed with Robetta’s RoseTTAFold (Baek, et al. 2021). Structures were visualized in PyMol (The PyMOL Molecular Graphics System, Version 2.0 Schrödinger, LLC). Secretion substrate prediction was performed with BastionX (Wang, et al. 2021) and EffectiveDB (Eichinger, et al. 2016). Dotplots were created in RStudio using the seqinr package, with parameters wsize = 20, wstep = 5, nmatch = 5.

### Insect rearing and developmental series

All insects were maintained at 25 °C on a 24-hour, 12:12 light:dark cycle. Wasp colonies included: *Trichogramma pretiosum “*Insectary line” infected with *Wolbachia* strain *w*Tpre (Lindsey, et al. 2016; Lindsey, et al. 2018a) and *Leptopilina clavipes* “LcNet” infected with *Wolbachia* strain *w*Lcla (Pannebakker *et al*, 2004; Kraaijeveld *et al*, 2016; Kampfraath *et al*, 2019; Lue *et al*, 2021). *Leptopilina clavipes* were hosted on *Drosophila virilis* (National Drosophila Species Stock Center SKU: 15010-1051.87), maintained on standard Bloomington cornmeal-agar medium (Nutri-fly® Bloomington Formulation) under density-controlled conditions. *Trichogramma pretiosum* were hosted on UV irradiated *Ephestia kuehniella* (Beneficial Insectary).

To generate a synchronized developmental series, adult *Trichogramma pretiosum* were given egg cards (*Ephestia kuehniella* eggs adhered to cardstock with double-sided tape) for two hours. Adult wasps were removed, and egg cards were held under standard rearing conditions until collection. Time points included: (1) 2-4 hours post-parasitism, during the embryo stage, (2) three days post-parasitism, at the prepupal stage, when host *Ephestia* eggs turned gray, (3) seven days post-parasitism, at the red-eye pupal stage when ovaries are maturing, and (4) adult females less than 24 hours post-eclosion that had not accessed fresh host eggs. Sampling times were based on published developmental studies of *Trichogramma* (Flanders 1937; Volkoff and Daumal 1994; Volkoff, et al. 1995; Jarjees and Merritt 2002), were collected in biological triplicate, and flash frozen by placement at –80 °C for further processing (see below). To prepare a synchronized developmental series of *Leptopilina clavipes*, one-day-old adult female wasps were given early second-instar *Drosophila virilis* larvae and observed for parasitism. Parasitized fly larvae were left to develop for (1) two hours (embryo), (2) fourteen days (early pupation), or (3) 18 days (red-eye). Adult females were collected for gene expression analyses within 24 hours of eclosion and prior to host access. Each stage was collected in biological triplicate as pools of three individuals, except for the embryo stage in which pools were composed of six individuals, to adjust for the small size. Samples were flash frozen and stored at –80 °C for further processing.

### Nucleotide Extractions, PCR, RT-PCR, and qRT-PCR

DNA was extracted from flash frozen samples with the Monarch® Genomic DNA Purification Kit (New England Biolabs), including the on-column RNase treatment. RNA was extracted from flash-frozen samples with the Monarch® Total RNA Miniprep Kit (New England Biolabs) including the DNase treatment step. The LunaScript® RT Master Mix Kit was used for making cDNA, following manufacturer’s instructions. PCR reactions were performed with Q5® Hot Start High-Fidelity 2X Master Mix (New England Biolabs) in 20 µl reactions and products were run on a 1% agarose gel alongside an appropriately sized ladder (either New England Biolabs 100 bp or 1kb DNA ladder) and stained post-electrophoresis with GelRed® (Biotium). *pif* expression was assessed with the Luna® Universal Two-Step RT-qPCR Master Mix (New England Biolabs) following manufacturer’s instructions, and normalization to *ftsZ* (*Wolbachia* housekeeping gene). All reactions were run in technical duplicate alongside a standard curve and negative controls (including no template and no reverse transcriptase controls) on a QuantStudio™ 3 Real-Time PCR System (Applied Biosystems™). All primer sequences are in Supplemental Table S2.

### Cloning and Yeast Transformation

Coding sequences of *pifA* and *pifB* were PCR amplified from *w*Tpre-infected wasps using modified forward primers to facilitate cloning into pENTR-D/TOPO (Invitrogen). These constructs were transformed into One Shot Top10 competent cells (Invitrogen) following manufacturer’s instructions (see Supplemental Table S2 for primer sequences) and plated on selective media. Entry vectors were verified first with restriction digests, followed by whole-plasmid sequencing (Plasmidsaurus). Validated entry vectors were recombined into an appropriate destination vector (see Supplemental Table S3) in an LR clonase reaction following manufacturer’s instructions (Invitrogen). Expression vectors were cloned into One Shot Top10 competent cells (Invitrogen), plated on selective media, and validated as above. Yeast strain W303 MATa (Supplemental Table S4) was transformed with expression vector(s) using the PEG/Lithium acetate method (Yeast Transformation Kit, MilliporeSigma)(Gietz and Woods 2002) and plated on non-inducing selective synthetic media (Sigma-Aldrich® Yeast Synthetic Drop-out Media Supplements: without leucine). In all cases, plasmid extractions from *E. coli* were performed with the Monarch® Plasmid Miniprep Kit (New England Biolabs).

Yeast transformants were grown overnight at 30 °C in selective non-inducing media with 2% glucose as a carbon source. Cells were harvested by centrifugation and washed twice with sterile water, followed by resuspension in water to an OD_600_ of 1.0. A five step, 10-fold dilution series was prepared for each transformant and 3 µl spots were plated on either selective inducing (2% galactose) or selective repressing (2% glucose) media. After spot plates dried, they were incubated at 30 °C for 24 hours.

### Microscopy and Image Analysis

For fluorescence imaging, yeast transformants were grown overnight at 30 °C in selective non-inducing media, supplemented with 2% raffinose as a carbon source, and 20µg/mL of additional adenine to minimize autofluorescence (Rines et al., 2011). Cells were harvested via centrifugation and resuspended in selective inducing media (2% galactose) prior to incubation at 30 °C for an additional six hours. After the 6-hour induction period, yeast cells were stained with NucBlue™ Live ReadyProbes™ Reagent (Hoechst 33342; Invitrogen™). Images were taken on an ECHO Revolve Fluorescence Microscope fitted with an 100x Apochromat Oil Immersion lens, FITC and DAPI LED filter sets, and an ELWD Universal Condenser. Images were merged and pseudocolored in FIJI (Schindelin, et al. 2012).

### Statistics and data visualization

Statistical analysis and data visualization were performed in R 4.1.3 (R Core Team, 2014). Variation in *pif* expression was assessed with a two-way ANOVA including gene and developmental stage as fixed effects, followed by post-hoc testing with the Tukey-Kramer method. Schematics were created with BioRender.com and data visualization leveraged the “muted qualitative colour scheme” designed by Paul Tol (https://personal.sron.nl/~pault/).

## RESULTS

### Comparative genomics identifies candidate parthenogenesis inducing factors (*pifs*)

We identified 10 sets of orthologous proteins (a total of 11 individual proteins from *w*Tpre and 20 from *w*Lcla) that were both (1) shared between *w*Lcla and *w*Tpre, and (2) absent from all other *Wolbachia* strains in our dataset. Twenty-seven of these genes (eight complete orthogroups) were pseudogenized (Supplemental Table S5). The remaining two sets of orthologous proteins were putatively functional, and each contained one hypothetical protein from *w*Lcla and one from *w*Tpre. For simplicity, these two sets of 1:1 orthologs are hereinafter referred to as PifA and PifB, for putative <u>P</u>arthenogenesis Inducing <u>F</u>actors A and B.

While the clustering of orthologous *Wolbachia* proteins identified only *w*Tpre and *w*Lcla as having PifA and PifB orthologs, we considered that (A) these proteins might be found in the published PI *Wolbachia* strains that we did not include in our database (*e.g*., *w*Uni (Gottlieb, et al. 2002), (B) there may be divergent Pifs present in other *Wolbachia* that did not meet clustering thresholds, (C) there might be unannotated *pif*-like regions present in other *Wolbachia* genomes, and, (D) there may be Pif-like proteins in other organisms. To address some of these issues, we used blast to search for putative Pif-homologs across non-redundant NCBI databases. Both protein and nucleic acid searches (blastp, tblastn, blastn, and discontiguous megablast) of *pifA* homologs only recovered matches to the already identified *w*Tpre and *w*Lcla PifA loci indicating these are unique genes restricted to these PI *Wolbachia* strains. Protein blast (blastp) searches of PifB identified short regions of distant homology to other proteins that contained leucine-rich repeat (LRR)-like domains. Outside of the *w*Tpre and *w*Lcla PifB orthologs, the highest scoring hit was a protein from an Alphaproteobacterium that aligned to 19% of the PifB sequence (at the LRR domains), with 30% amino acid identity (Supplemental Figure S1, Supplemental Table S10). Lower scoring hits included a small number of proteins from other *Wolbachia* that had also acquired LRR-like domains, but, like the other database matches, they did not appear to be homologs of the *w*Tpre and *w*Lcla PifB proteins. Regions of similarity were divergent and restricted to the LRR-like domains, and the proteins did not globally align to the *w*Tpre and *w*Lcla Pifs or have any nucleotide-level sequence homology (Supplemental Figure S1, Supplemental Table S10).

Both PifA and PifB have canonical features of eukaryotic host-associated bacterial effector proteins, including the aforementioned restricted phylogenetic patterns. Additionally, there are several shared characteristics with *Wolbachia* proteins that mediate other reproductive phenotypes. These similarities include proximity to phage and other mobile element loci, and location within a putative eukaryotic association module (EAM) (Figure 1C). The EAM is a region found in many *Wolbachia* genomes: it is associated with *Wolbachia* prophage (WO) regions or remnants, and in other *Wolbachia* strains the EAM contains loci responsible for cytoplasmic incompatibility and/or male killing (Bordenstein and Bordenstein 2016; LePage, et al. 2017; Lindsey, et al. 2018b; Perlmutter, et al. 2019). *Trichogramma*-infecting *Wolbachia* were previously identified as having lost their WO phages (Gavotte, et al. 2007), and genome sequencing of the *w*Tpre strain revealed only degenerate prophage WO regions (Lindsey *et al*, 2016). Indeed, the *pif* loci in *w*Tpre are surrounded by those degenerate phage and transposable element loci, as well as other pseudogenes consistently found in the EAM, including *mutL*, a patatin-like gene, transcriptional regulators, and a suite of ankyrin domain containing proteins (Figure 1C). The homologous *pif* loci in *w*Lcla are contained within a short contig (wLcla_Contig_2; ∼11kb) that also includes transposases, a transcriptional regulator, and pseudogenized hypothetical proteins. Due to the fragmented nature of the *w*Lcla assembly, it is difficult to determine if the *w*Lcla *pif* homologs are in a true prophage and/or EAM-like region. Finally, while *pifA* and *pifB* are syntenic in *w*Tpre (329 bp between the 3’ end of *pifA* and the 5’ end of *pifB*), in *w*Lcla the two genes are in opposite orientations and separated by a ∼1.6kb region that contains a pseudogenized transposase and two small hypothetical proteins.

PifA and PifB contain eukaryotic-like domain structures: another key indicator of bacterial proteins that manipulate eukaryotic biology. Structural prediction revealed four regions of the Pif proteins with significant structural homology to eukaryotic domains (Figure 2A, Supplemental Tables S6-S9). The first is a domain with high similarity to nucleoporins at the N-terminus of PifA. Additionally, PifA contains two overlapping domains predicted to function in RNA splicing. Most notably, the second of these has significant structural similarity (33%) to the insect protein Transformer (Tra), discussed above for its role as the master-regulatory sex determination gene (Verhulst *et al*, 2010). Finally, PifB is primarily composed of a large leucine-rich repeat (LRR) domain with strong structural homology to eukaryotic LRR receptor-like serine/threonine-protein kinases, especially membrane-bound Toll-like receptors (Figure 2A). *w*Tpre homologs of the Pif proteins were modeled with Robetta to visualize putative 3D protein structures (Figures 2B-C, Supplemental Files S1-S4). The predicted PifA protein structure consisted of a pore-like formation in the nucleoporin-like region (error estimate <10) and a relatively unstructured region aligning with the predicted RNA splicing/Tra-like domain. PifB structure results produced a double coiled structure typical of LRRs. Lastly, PifA and PifB are predicted to be bacterial secretion substrates, with a strong likelihood of secretion via the Type IV Secretion System (T4SS), which is known to translocate *Wolbachia* effector proteins to the host in other strains (Whitaker, et al. 2016)(Supplemental File S11).

**Figure 2.**
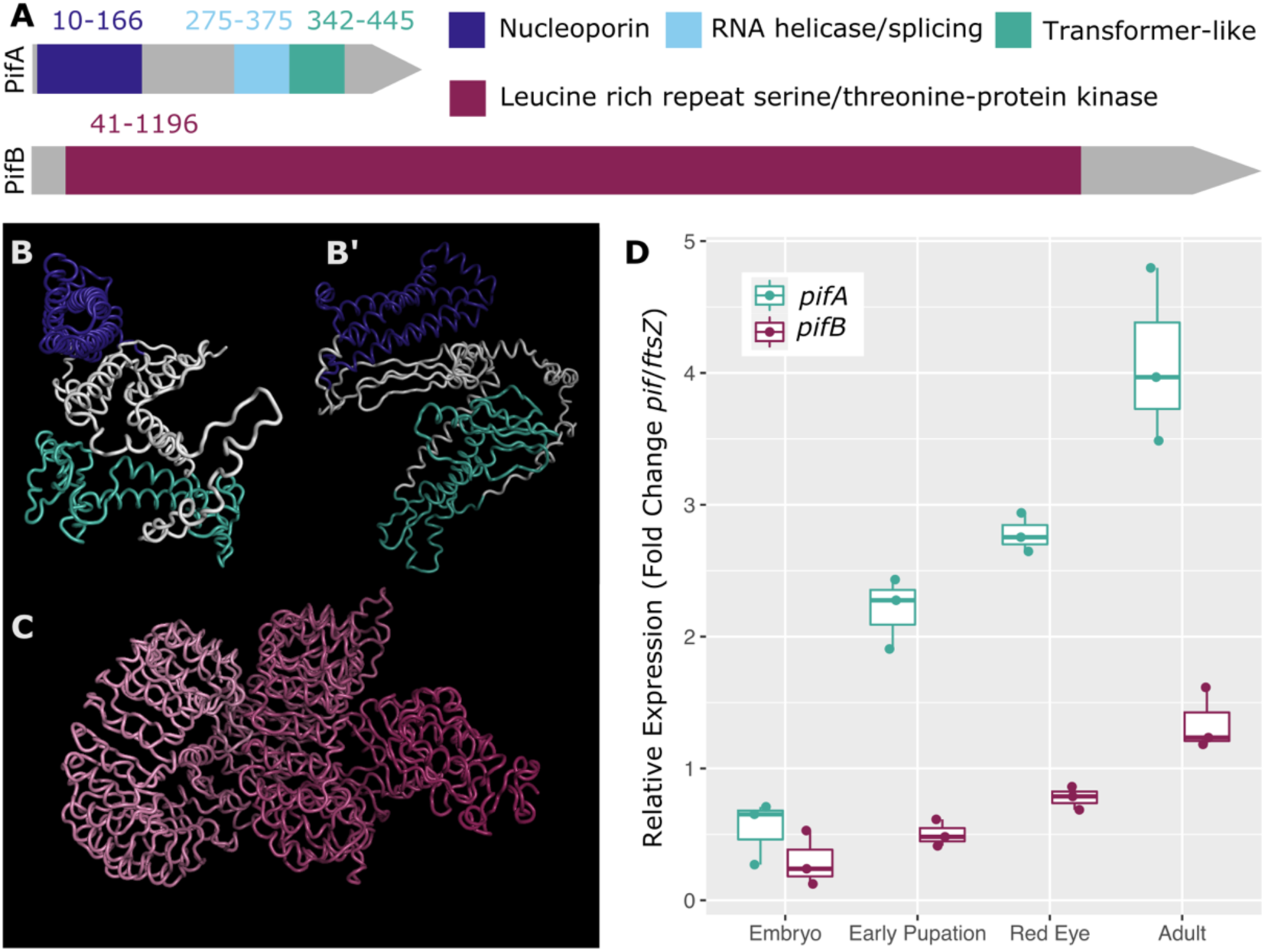
Candidate parthenogenesis-inducing factors PifA and PifB. (**A**) Domain prediction indicates PifA and PifB contain eukaryotic-like structures. Colored coordinates indicate amino acid positions of the relevant domain in the *w*Tpre ortholog. **(B and B’)** Two views of the predicted structure of the PifA protein from *w*Tpre. B’ view is rotated 90 degrees clockwise around the vertical axis as compared to the view in (B). The nucleoporin-like domain is in indigo, and the RNA processing-like region encompassing both the RNA helicase and Tra-like regions is in teal. **(C)** Predicted structure of the PifB protein from *w*Tpre. The color gradient is for ease of viewing the 3D structure. **(D)** *pif* expression (relative to a bacterial house-keeping gene) increases across wasp development in *Trichogramma pretiosum*.

### *Wolbachia* infection is correlated with differential splicing of *transformer* and *double-sex*

Given the discovery of a Tra-like *Wolbachia* protein, and data indicating that *Wolbachia* strain *w*Lcla impacts sex-specific splicing of *tra* in *L. clavipes* females (Geuverink, et al. 2018), we queried previously published RNA-seq data from adult female *Trichogramma pretiosum* to assess the impacts of *Wolbachia* on *tra* and *dsx* splicing. These data are derived from three sets of wasp colonies, each with a genetically identical, *Wolbachia*-infected and –uninfected pair (n=6 total). Exon use was previously assessed genome-wide, and here we display these patterns specifically for *tra* and *dsx* (Figure 1D-E). We find that in female wasps without *Wolbachia,* exon five of *tra,* and exons 3-4 of *dsx* are uniquely excluded. However, while female wasps with *Wolbachia* do use *tra* exon 5, there was no identifiable expression of *tra* exons 1-2.

### *pif* expression correlates with insect reproduction

To see if *pifA* and *pifB* are expressed, and thus less likely to be an artifact of genome annotation, we used RT-PCR to amplify *pifA* and *pifB* transcripts from a pool of ∼2-4-hour old *Trichogramma pretiosum* embryos, and a pool of adult females. Indeed, both *pifA* and *pifB* are expressed in adult females, and we detected low or no expression of *pifA* and *pifB* in the young embryos (Supplemental Figure S2). Because *pifA* and *pifB* are syntenic in *w*Tpre, reminiscent of the cytoplasmic incompatibility loci which are co-transcribed (Lindsey *et al*, 2018), we also used RT-PCR to look for the presence of *pifA-pifB* monocistronic transcripts, indicative of an operon. We were not able to amplify any monocistronic transcripts from adult females using primers that targeted the 5’ region of *pifA* and the 3’ region of *pifB* (Supplemental Figure S3).

To determine if *pif* expression correlated with reproductive biology, we used qRT-PCR to quantify relative *pifA* and *pifB* expression across four time points in *T. pretiosum* development: early in embryogenesis, during the transition to metamorphosis, the “red eye stage” of metamorphosis during which gonads undergo differentiation, and in adult females (Figure 2D). There was a statistically significant increase in both *pifA* and *pifB* expression across wasp development (F_1,3_= 61.4, p < 0.0001). However, the relative expression of *pifA* was always significantly higher than *pifB* post embryonic development (F_1,3_=18.51, p < 0.0001). At 2-4 hours post-parasitization, (shortly after the first mitotic division during which the *Wolbachia*-mediated gamete duplication step occurs (Pannebakker, et al. 2004), there was minimal expression of *pif* genes. In contrast, in freshly emerged adult females, expression of *pifA* and *pifB* was 4– and 1.5-fold higher than in embryos, respectively. Expression patterns of *pifA* and *pifB* in *w*Lcla and *L. clavipes* recapitulated these patterns, though *pifA* had even greater expression than *pifB* (up to 25X higher) (Supplementary Figure S4).

### PifA is conserved and repetitive in the Tra-like region

In validating plasmids and sequencing entry amplicons (see methods), we discovered that the *w*Tpre *pifA* gene contains a short region present in tandem triplicate that had been collapsed into a duplicate during genome assembly. The *w*Lcla PifA ortholog was assembled correctly, based on our amplicon sequencing, though there were similar repeated regions in the same part of the coding sequence. Given these findings on the repetitive nature of PifA, we next characterized these patterns and their conservation. First, we compared the two orthologs to each other and we found a region of high similarity aligning with the Tra-like domain (Figure 3A). Here, there is a ∼100 amino acid region of high synteny and conservation. This conserved region itself composed of tandemly repeated blocks of 12-18 amino acids (seen in the classic clustering of the dot plot in that region; Figure 3A). To better visualize these tandem repeats, we compared each ortholog to itself (Figure 3B-C). In *w*Tpre, there are 4 of repeated segments in total, each 18 amino acids long (Figure 3B). In *w*Lcla the repeated region was composed of 6 amino acid segments, 12 long, and there was an additional instance of that repeated segment in the ∼250 amino acid region (Figure 3C).

**Figure 3:**
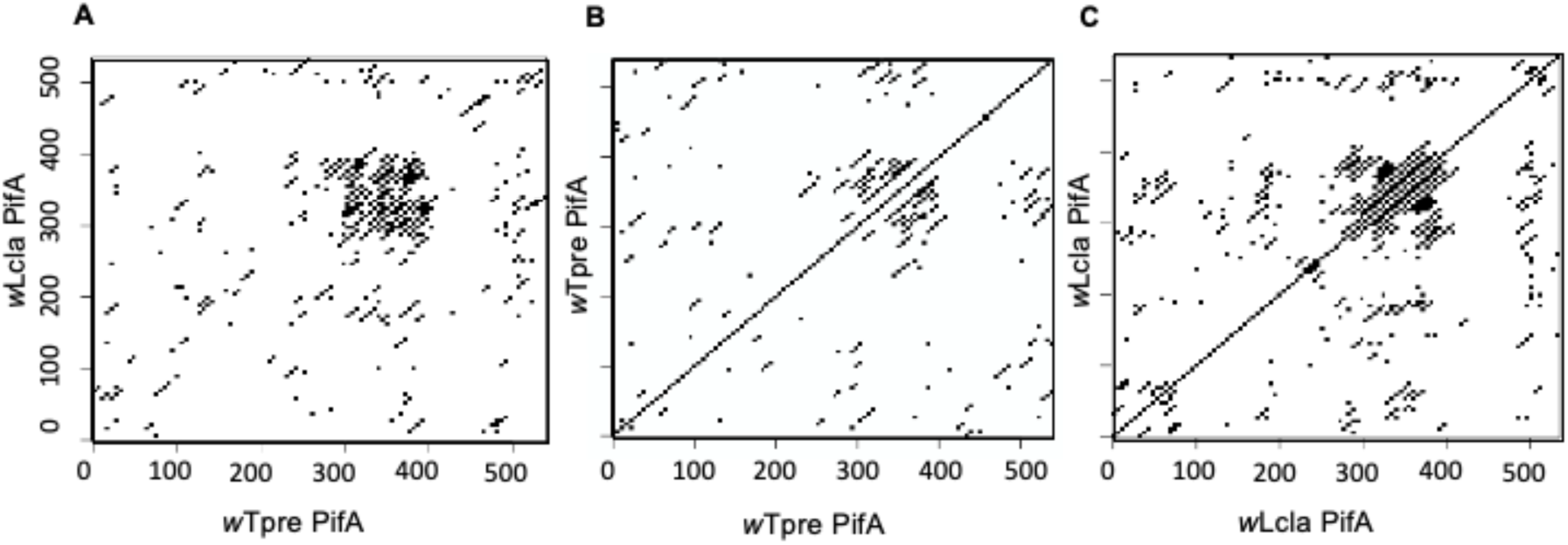
PifA is conserved and repetitive in the Tra-like region. Dot-plots showing amino acid similarity, where each point represents a window of five identical amino acids. (**A**) *w*Tpre PifA versus *w*Lcla PifA, showing high conservation between the orthologs in the ∼320-420 amino acid region. **(B)** *w*Tpre PifA versus self, showing a series of repeated regions in the Tra-like region. **(C)** *w*Lcla PifA versus self, showing a series of repeated regions in the Tra-like region.

### PifA and PifB impact eukaryotic cells

To determine if PifA and PifB are putative effector proteins, we used a standard screen in the budding yeast, *Saccharomyces cerevisiae*. Bacterial effector proteins that interact with eukaryotic cell biology often cause growth defects when expressed in yeast cells (Kramer, et al. 2007; Siggers and Lesser 2008; Sheehan, et al. 2016; Beckmann, et al. 2017; Rice, et al. 2017). We expressed the *w*Tpre PifA and/or PifB genes in W303 MATa yeast using a 2-micron expression vector with a galactose-inducible expression system, pAG424GAL-ccdB and/or pAG425GAL-ccdB (Alberti, et al. 2007)(Figure 4). At 24 hours, there were identifiable growth defects in the yeast expressing PifA or PifB, relative to the empty vector control. Co-expression of both Pifs also caused growth defects relative to the double vector control. We note that the yeast transformed with two high copy number 2-micron expression vectors generally do not grow as well, but there are still clear differences between the empty and Pif-containing conditions. By 48 hours, there was still evidence for negative impacts on yeast growth in the PifA and co-expressing yeast. However, the yeast expressing only PifB were more similar to the empty vector controls at this later timepoint.

**Figure 4:**
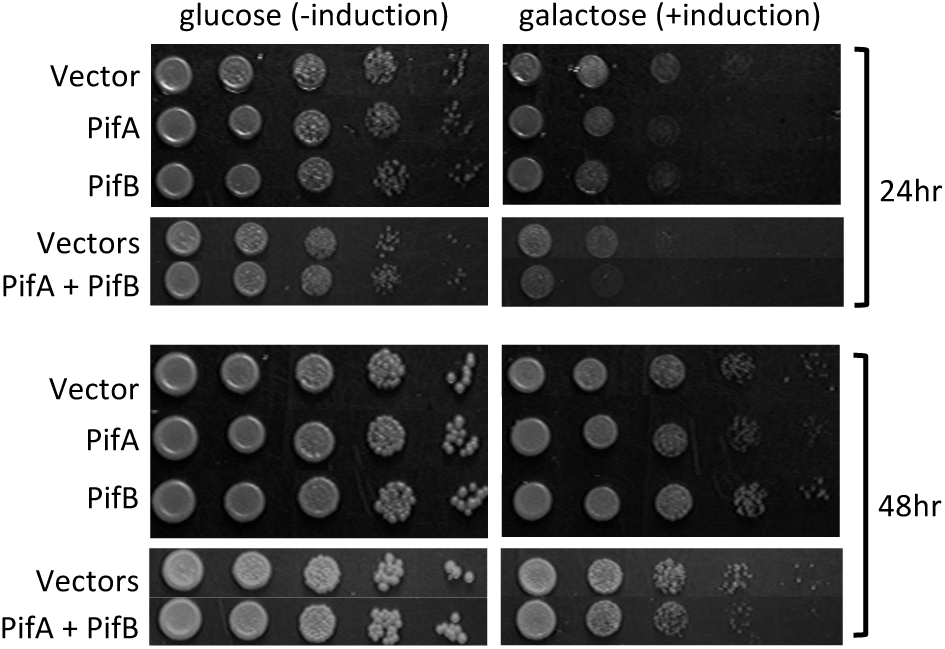
PifA and PifB impact eukaryotic cell biology. PifA and PifB cause growth defects in yeast relative to empty-vector and no induction controls. Yeast strain W303 MATa was transformed with galactose-inducible 2-micron expression vectors that were either empty or contained a Pif CDS. Transformants were grown up in selective non-inducing media, normalized to OD600 = 1, and serial 1:10 dilutions were spotted on selective inducing or non-inducing plates and grown at 30 °C for 48 hours. Single vector conditions leveraged the pAG425 backbone, and double expression conditions leveraged both pAG424 and pAG425 (see supplemental materials).

### PifA localizes with atypically dividing nuclei

Finally, given the stronger growth impacts of PifA, and its nucleoporin-like domain, we used a fluorescent fusion tag to localize PifA within the yeast cells. GFP alone displayed a homogeneous localization pattern spread evenly throughout the yeast cytoplasm (Figure 5A). In contrast, GFP-PifA exhibited strong nuclear or perinuclear localization, reminiscent of localization patterns of known nucleolar proteins (Carmo-Fonseca, et al. 2000; Taddei and Gasser 2012)(Figure 5B-D). Some yeast cells exhibited GFP-PifA puncti near each pole of dividing nuclei and deposition into the daughter cell (Figure 5B,C). Other yeast cells had what looked like PifA dispersed around the dividing nucleus (Figure 5D). Importantly, there was also evidence for atypical mitoses in the presence of PifA (Figure 5D) with uneven distribution of daughter nuclei in the two cells.

**Figure 5:**
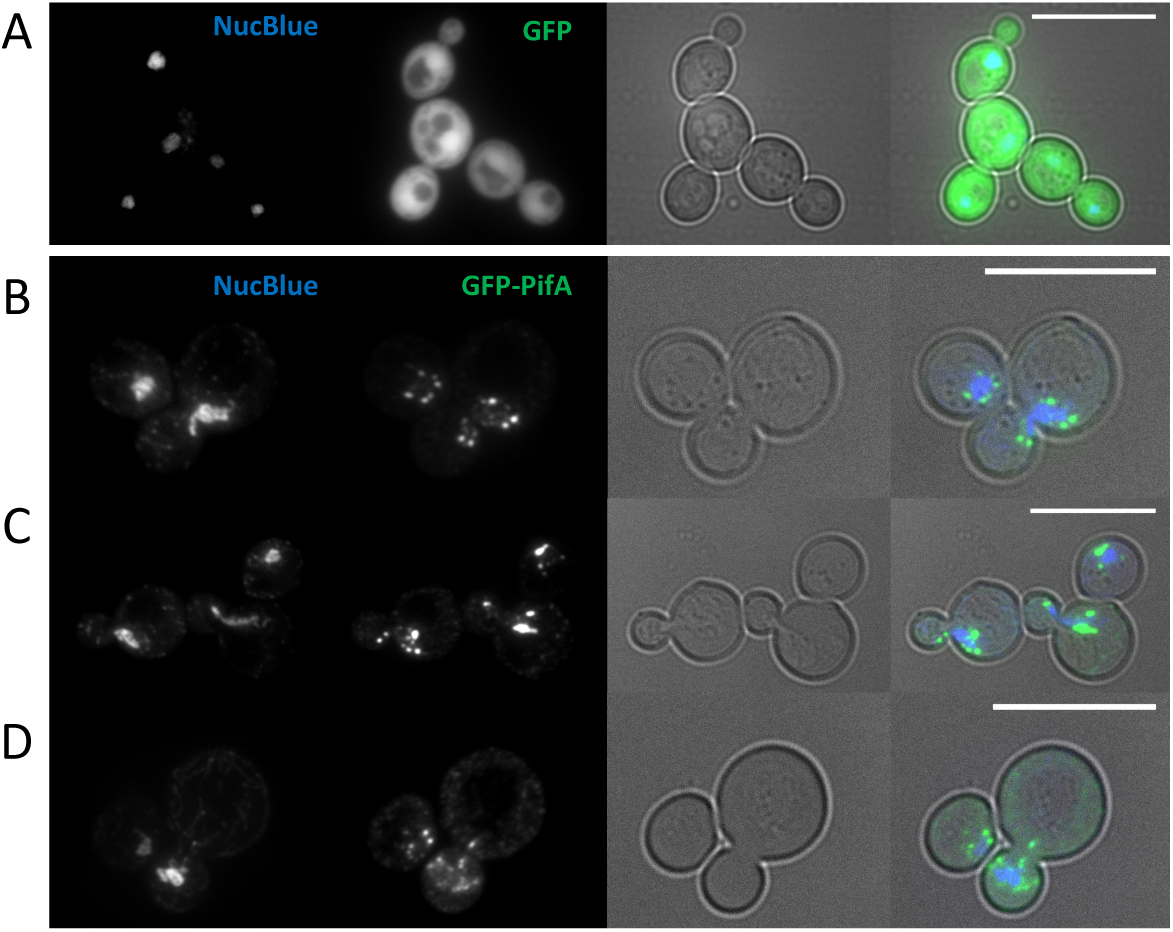
PifA associates with dividing nuclei. To localize PifA proteins, yeast W303 MATa were transformed with galactose-inducible pFUS expression vectors (with an N-terminal GFP tag) and grown in selective inducing media (with 2% galactose as a carbon source) for six hours, and stained with NucBlue. Scale bars represent 10 microns. **(A)** Yeast with empty pFus, expressing GFP alone. **(B-D)** Yeast with pFus-PifA, expressing GFP-PifA.

## DISCUSSION

For decades the genes responsible for PI have remained elusive. This is despite increasing examples of microbe-mediated PI in nature, and significant progress made in understanding the mechanisms of other microbial reproductive manipulations such as cytoplasmic incompatibility and male-killing (Kaur et al., 2021). Indeed, PI is present in *Wolbachia* strains infecting at least six separate genera of parasitoid wasps (Ma and Schwander 2017). Because PI evolved multiple times *across Wolbachia*, we focused on strains predicted to share the same two-step mechanism of PI. As such, we identified and characterized two *Wolbachia* proteins that we predict are involved in the induction of parthenogenesis in *Trichogramma pretiosum* and *Leptopilina clavipes:* PifA and PifB.

Broadly, the proteins we identified have many characteristics of host-associated bacterial effector proteins. These include Pif-mediated growth phenotypes in yeast and predictions that the Pifs are Type IV Secretion substrates: bacterial protein translocation systems that both *w*Lcla and *w*Tpre encode for (Lindsey 2020). We do note that, yeast and insect mitoses have a number of key differences (e.g., open versus closed, (Sazer, et al. 2014), and any signal indicating egg fertilization would likely not manifest in yeast cells, so it is unsurprising that the growth impacts in yeast are more mild than many other pathogenic bacterial effector proteins with broad host ranges (e.g., in *Legionella* (Siggers and Lesser 2008)).

Regardless, the predicted domain structures of the Pif proteins are highly characteristic of functions in eukaryotic cells. PifA has both a nucleoporin-like region, and an RNA-splicing-like region with homology to the insect-specific protein Transformer with is a strong indicator of interaction with host sex determination. The structural homologies of PifB to serine/threonine kinases, leucine-rich repeats (LRR), F-box, and Toll-like receptor motifs indicate the ability to recognize, bind, and/or alter other proteins. LRR motifs are widespread in eukaryotic proteins and are typically involved in mediating protein-protein interactions (Kobe and Kajava 2001). Consequently, LRR motifs are also found in other bacterial effector proteins, acting to mimic and interface with host biology (Aepfelbacher, et al. 2005; Xu, et al. 2008).

In line with our hypotheses about the diversity of mechanisms underlying PI, there is no evidence for these proteins in other published *Wolbachia* genomes, including strains that use other PI mechanisms, based on blast results. Results from computational predictions for PifA and PifB also align with our expectation that two-step PI *Wolbachia* need machinery for interacting with both chromosome division, and the sex determination system. In two-step PI systems, the diploidization event during early embryogenesis is critical. This diploidization (in *w*Tpre and *w*Lcla, the failed anaphase) appears to be highly regulated as it only ever occurs during the first embryonic mitosis. If diploidization does not occur at the first mitosis (perhaps due to reduced *Wolbachia* titers), then it does not appear to happen at all (Tulgetske 2010). Additionally, PI *Wolbachia* do not appear to cause rampant polyploidy, and, *Wolbachia*-mediated diploidization does not occur in infected embryos if they have been fertilized (Stouthamer and Kazmer 1994b). In many *Trichogramma,* PI-*Wolbachia* infected females can mate and fertilize their eggs, in which case, the resulting offspring are diploid females, and not triploid offspring (Stouthamer and Kazmer, 1994), indicating that the PI mechanism is sensitive to both the specific mitotic event, and to whether the egg has been fertilized.

Considering the specific conditions in which diploidization takes place, we hypothesize there is a host factor that serves as a signal for both the timing of the diploidization event (the first mitosis), and if diploidization is “not needed” (i.e., because the egg was fertilized). We argue it is unlikely for embryo ploidy itself to directly serve as this signal, for several reasons. First, in insects, maternal and paternal pronuclei stay physically separated for most of the first mitotic cycle (Tram, et al. 2003). Second, even if *Wolbachia* or PI factors were capable of “counting chromosomes”, a ploidy signal would not explain why the diploidization event only ever occurs at the first mitosis. Finally, interactions with a separate male sex-ratio distorter that is present in some *Trichogramma* species, the “Paternal Sex Ratio” (PSR) chromosome, indicate *Wolbachia*-mediated PI is sensitive to fertilization independent of ploidy. Specifically, the PSR “B-chromosome” is transmitted by haploid males: upon fertilization, the PSR chromosome mediates destruction of all other paternal chromosomes, resulting in a haploid male (a single maternal genome copy) plus the PSR chromosome which can then be transmitted to the next generation. Critically, when these PSR-males mate with *Wolbachia*-infected females, PSR-mediated distortions take precedence over *Wolbachia*-mediated PI (Stouthamer, et al. 2001). The paternal chromosomes are destroyed, leaving a haploid embryo with PSR, but *Wolbachia-*mediated diploidization never occurs. Thus, there is likely a non-chromosomal factor associated with fertilization, or its absence, that ultimately makes *Wolbachia*-mediated PI in this system possible both (1) during a specific mitotic event and (2) in the absence of fertilization.

A potential non-chromosomal factor that could signal both of these conditions is the presence or absence of paternally contributed centrioles. In fertilized hymenopteran embryos (e.g., *Nasonia vitripenis*), the sperm-tail basal body preferentially nucleates paternal centrioles to take part in the first mitotic division, and maternal cytoplasmic asters disperse (Ferree, et al. 2008). In unfertilized, haploid embryos, this process is compensated for by *de novo* centrosome formation in the embryo from the maternal cytoplasmic asters (Ferree, et al. 2006). The absence of a paternal centrosome in unfertilized eggs may serve as a signal or is possibly functionally significant in the PI mechanism. In the unfertilized embryo, if the PI proteins interact with the *de novo* formation of centrosomes, inhibition of proper chromosome segregation may result. Indeed, the centrosome looks to be a target of other sex-ratio modifying infections such as the male-killing bacterium *Arsenophonus nasoniae* in *Nasonia vitripennis*. Here, the bacterium suppresses maternal centrosome formation in the unfertilized, male destined embryos. This results in mitotic spindle disruption, failed development of the early embryo, and ultimately male-specific mortality (Ferree et al., 2008).

Regardless of what the specific target of PI *Wolbachia*-mediated diploidization may be, we do know that an extensive network of protein kinases is required for successful mitosis in animal cells. These protein kinases, such as Aurora A, Aurora B, and polo kinase in *Drosophila melanogaster*, have multiple roles in regulating spindle assembly, centrosomal functioning, and cytokinesis (Archambault and Carmena 2012; Reinhardt and Yaffe 2013). The serine/threonine protein kinase-like domain of PifB may indicate participation in these mitotic processes. Additionally, in many eukaryotes, mitotic phosphorylation sites typically lie in serine/threonine motifs (Canova and Molle 2014). Alternatively, PifA may be participating in mitotic disruption, as the N-terminal region has structural homology to a suite of nucleoporins that drive spindle assembly and chromosome organization in wide range of other organisms (Chatel and Fahrenkrog 2011). Indeed, many of the PifA localization patterns in dividing yeast cells (Figure 5) are reminiscent of the nucleoporin ALADIN in *Drosophila melanogaster* (Carvalhal, et al. 2015). ALADIN aggregates at the poles of dividing nuclei, and regulates localization of Aurora A, a serine/threonine kinase and spindle organization (Carvalhal, et al. 2015). There is also potential for two proteins to interact to drive the failed first anaphase. For example, one protein’s activity might be sensitive to the presence of the critical host factor (e.g., perhaps centrioles, as discussed above), and the other protein could then recruit the first protein to the critical host structures. This is of course speculation, but the localization patterns of PifA and mitotic impacts hint at more than interaction with the sex determination pathway.

In addition to this tightly regulated diploidization event, a second step is required to achieve female development in the *w*Lcla and *w*Tpre PI systems. In hymenopterans the proper initiation of female development relies on a paternally derived signal. This might be in combination with a maternal effect (as in *Nasonia vitripenis* (Beukeboom and Van De Zande 2010)), or the paternal signal might be sufficient on its own (as is predicted to be the case in *Leptopilina clavipes* (Chen 2021)). In *Nasonia vitripennis*, the maternal *tra* allele is transcriptionally repressed in the embryo, and a combination of maternally loaded *traF* mRNAs and a transcriptionally active paternal *tra* allele is necessary for female development (Beukeboom and Van De Zande 2010). In contrast, maternally provided *traF* is absent in *Wolbachia-*uninfected *Leptopilina clavipes* embryos (Chen 2021), and it may be the case that paternal activation of *traF* is sufficient. Importantly, *tra* isoforms differ between *Wolbachia*-infected and *Wolbachia*-uninfected adult female *Leptopilina clavipes*. In the *Wolbachia*-infected wasps, only the *traF* variant is present (Geuverink, et al. 2018). In the *Wolbachia*-uninfected wasps, both *traF* and *traM* splice variants are present: perhaps due to somatic *traF* and *traM* in developed oocytes. Regardless of maternal effect, the absence of paternally derived *traF* during asexual reproduction presents a constraint on female development and an explanation for *Wolbachia*’s interference with the sex determination cascade.

The Tra-like domain within PifA may function as a replacement to initiate *tra* autoregulation and/or act on downstream sex determination genes such as *dsx* to ensure female differentiation. The nuclear localization of PifA in yeast cells notably complements its predicted nucleoporin-like region, suggesting that it may function within the host nucleus, or perhaps on the nuclear membrane where post-transcriptional processing often occurs (Keene 2007). Indeed, previous experiments in *Trichogramma pretiosum* found that *Wolbachia* infection results in a number of differentially expressed and differentially spliced transcripts, and included in these are genes in the sex determination pathway (Wu, et al. 2020)(Figure 1D,E). Given the predicted structure of PifA, these patterns could be attributed to direct splicing of host RNAs. However, it is also possible that PifA may act via allosteric or competitive interactions with the spliceosome or other RNA binding proteins.

PI microbes are clearly rich in factors for manipulating chromosome biology and sex determination of their hosts. Considering the impact of PifA and PifB on yeast biology, their location in a eukaryotic-association module genomic region, and structural similarities to eukaryotic domains involved in mitosis and sex determination, these proteins are promising candidates for Parthenogenesis Induction. At present, we are quite limited by the lack of genetic tools in *Wolbachia* and natural hosts of PI-*Wolbachia*. Investigation of other *Wolbachia* phenotypes such as CI and male killing have leveraged accessible heterologous expression systems in the *Drosophila melanogaster* model: an insect that naturally harbors such symbionts. Indeed, tools in parasitic hymenopterans or other haplodiploid arthropods are quite sparse. Even in the most well-studied species, *Nasonia vitripenis,* CRISPR has only recently been used for gene-editing (Li, et al. 2017; Chaverra-Rodriguez, et al. 2020), but as of yet no one has achieved integration of a transgene in any Hymenopteran. Ongoing investigation into the mechanisms of symbiont-mediated PI and deciphering the involvement of PifA and PifB will thus require a stronger focus on biochemical approaches and methods development for cell biology and genetics in these difficult to work with insects. Regardless, our study represents the first identification of putative parthenogenesis-inducing proteins, a major step towards understanding host-symbiont interactions and the evolution of asexual reproduction.

## DECLARATIONS

## Acknowledgements

Research reported in this publication was supported by the National Institute of General Medical Sciences of the National Institutes of Health under award number R35GM150991. This work was also supported by UMN AGREETT startup funds, a UMN New Faculty Grant awarded to ARIL, and a USDA Predoctoral Fellowship to LCF (NIFA 2023-67034-40496). LCF was additionally supported by UMN DOVE and UMN CFANS Match fellowships. We thank Danny Rice for feedback on genomic data, Irene Newton for gifting us pFUS and yeast strain W303, and Nathan Mortimer for stocks of *Leptopilina clavipes* and *Drosophila virilis*.

## SUPPLEMENT

### Supplemental Excel File Contains

Table S1. Genomes used in comparative analyses

Table S2. Primers

Table S3. Vectors

Table S4. Yeast strains

Table S5. Orthogroups unique to *w*Tpre and *w*Lcla

Table S6. HHpred results for *w*Tpre PifA

Table S7. HHpred results for *w*Tpre PifB

Table S8. HHpred results for *w*Lcla PifA

Table S9. HHpred results for *w*Lcla PifB

Table S10. PifB BLASTP Results

Table S11. Secretion prediction results for wLcla and wTpre PifA and PifB homologs

### Supplemental File S1 Contains

#### Supplemental Methods

Figure S1. PifB BLASTP matches outside of *w*Tpre and *w*Lcla are restricted to low similarity LRR-like domains in other bacteria.

Figure S2. *pifA* and *pifB* are expressed in adult female *Trichogramma pretiosum*.

Figure S3. *pifA* and *pifB* are not co-transcribed in *Trichogramma pretiosum*.

Figure S4. Time series of *w*Lcla *pifA* and *pifB* expression in *Leptopilina clavipes*.

#### Other Supplemental Files

Supplemental File S2: Robetta results for PifA

Supplemental File S3: Robetta results for PifB with 29 N-terminal amino acids deleted

Supplemental File S4: Robetta results for PifA with 29 C-terminal amino acids deleted

