## Supplemental Figures for "Identification of parthenogenesis-inducing effector proteins in *Wolbachia*"

### Supplemental Figures for: Identifying parthenogenesis-inducing effector proteins in *Wolbachia*

Laura C Fricke[1] and Amelia RI Lindsey[1*]
[1]Department of Entomology, University of Minnesota, St. Paul, Minnesota, 55108

#### Supplemental Figures

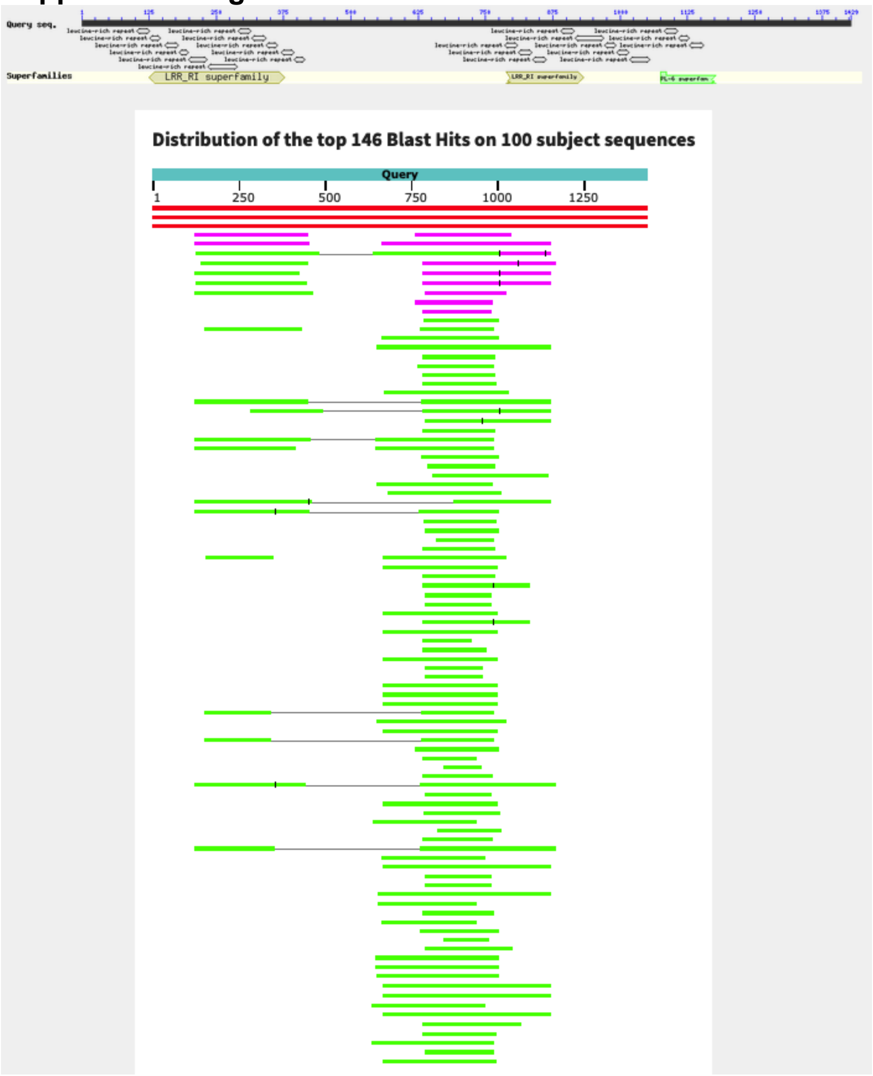


**Figure S1. PifB BLASTP matches outside of *w*Tpre and *w*Lcla are restricted to low similarity LRR-like domains in other proteins.**

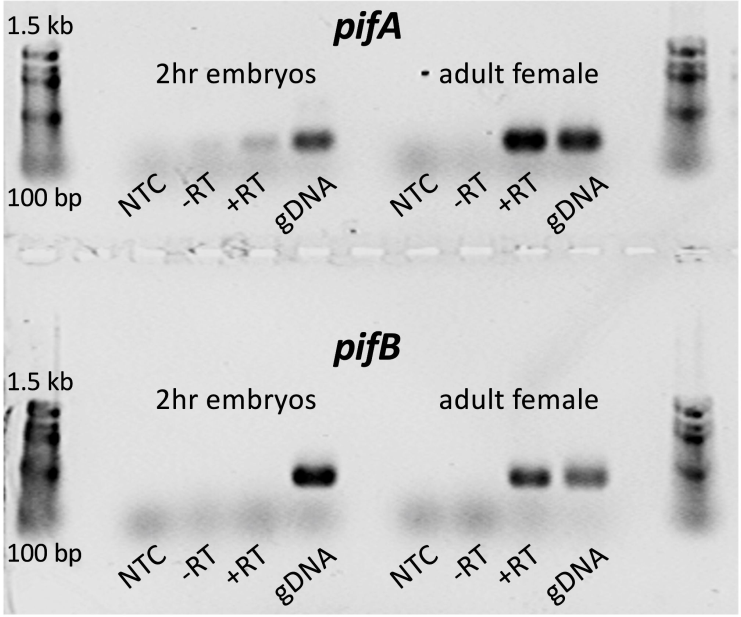


**Figure S2. *pifA* and *pifB* are expressed in adult female *Trichogramma pretiosum*.** RT-PCR was used to amplify ~300 bp regions of *pifA* and *pifB* from two hour old embryos, and adult females. Controls include no template (NTC) and no reverse transcription (-RT) negative controls, and a gDNA positive control.

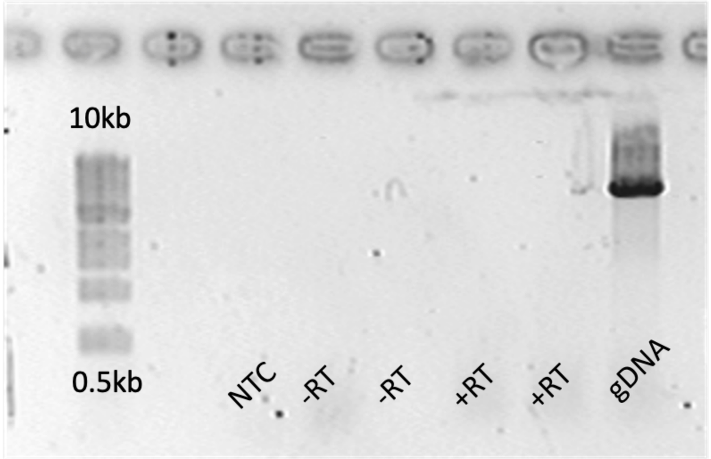


**Figure S3. *pifA* and *pifB* are not cotranscribed in *Trichogramma pretiosum.*** RT-PCR was used to amplify across *pifA* and *pifB* cDNA from adult females. Controls include no template (NTC) and no reverse transcription (-RT) negative controls, and a gDNA positive control.

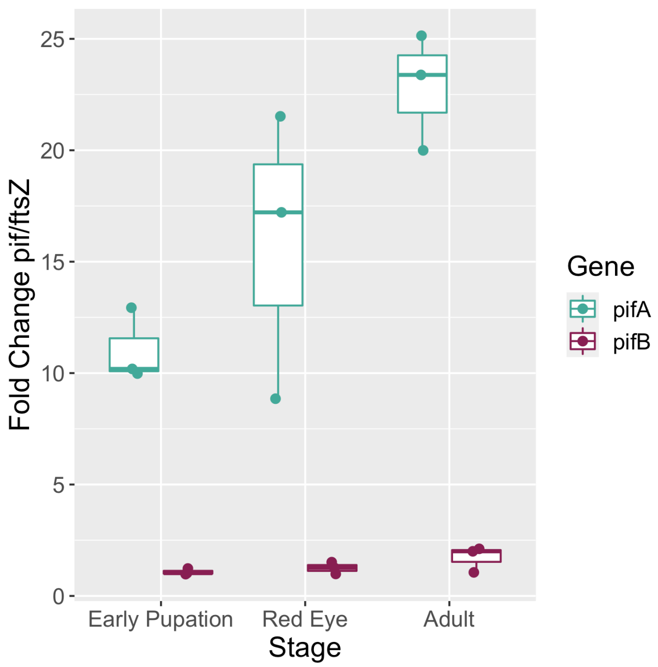


**Figure S4. *w*LcIa pif expression increases across wasp development**
qRT-PCR was used to quantify expression of *pif* loci across development, and was normalized to *ftsZ*. *pif* expression increases through wasp development for both *pifA* and *pifB*. There was no expression present in the early embryo.
